# Intersection of structural and functional connectivity of the nucleus basalis of Meynert in Parkinson’s disease dementia and Lewy body dementia

**DOI:** 10.1101/2020.07.27.221853

**Authors:** Ashwini Oswal, James Gratwicke, Harith Akram, Marjan Jahanshahi, Laszlo Zaborszky, Peter Brown, Marwan Hariz, Ludvic Zrinzo, Tom Foltynie, Vladimir Litvak

## Abstract

Parkinson’s disease dementia (PDD) and dementia with Lewy bodies (DLB) are related conditions that are associated with cholinergic system dysfunction. Dysfunction of the nucleus basalis of Meynert (NBM), a basal forebrain structure that provides the dominant source of cortical cholinergic innervation, has been implicated in the pathogenesis of both PDD and DLB. Here we leverage the temporal resolution of magnetoencephalography (MEG) with the spatial resolution of MRI tractography in order to explore the intersection of *functional* and *structural* connectivity of the NBM in a unique cohort of PDD and DLB patients undergoing Deep Brain Stimulation (DBS) of this structure. We observe that NBM-cortical *structural* and *functional* connectivity correlate within spatially and spectrally segregated networks including: 1) a beta band network to supplementary motor area (SMA), where activity in the SMA was found to drive activity in the NBM, 2) a delta/theta band network to medial temporal lobe structures encompassing the parahippocampal gyrus and 3) a delta/theta band network to visual areas including lingual gyrus. These findings reveal functional networks of the NBM that are likely to subserve important roles in motor control, memory and visual function respectively. Furthermore, they motivate future studies aimed at disentangling network contribution to disease phenotype.

## Introduction

Parkinson’s disease dementia (PDD) and dementia with Lewy bodies (DLB) are two of the commonest neurodegenerative dementias (Aarsland and Kurz, 2010; Jellinger, 2018). They are both characterised by the neuropathological hallmark of cortical Lewy bodies composed of alpha-synuclein (Hurtig *et al.*, 2000; Horvath *et al.*, 2013) and are associated with marked cholinergic neurotransmitter system dysfunction (Shimada *et al.*, 2009). They also share a common phenotype - including prominent executive, attentional and visual processing dysfunction, memory deficits, cognitive fluctuations, visual hallucinations and parkinsonism (Emre *et al.*, 2007; McKeith *et al.*, 2017). Clinical differentiation between these two interrelated conditions depends on whether dementia occurs in the context of established Parkinson’s disease (PDD) or prior to/concurrent with parkinsonian symptoms (DLB).

The nucleus basalis of Meynert (NBM) is the principle source of cholinergic inputs to cortex, which are implicated in memory, attention, visual processing and motor plasticity (Mesulam and Geula, 1988; Gratwicke *et al.*, 2013). Furthermore, degeneration of the NBM has been shown to predict cognitive impairments in both PDD and DLB (Whitehouse *et al.*, 1983; Choi *et al.*, 2012; Grothe *et al.*, 2014; Ray *et al.*, 2018). Strategies aimed at modulating the activity of the NBM and its cortical efferents have been proposed as therapies for both conditions (Gratwicke *et al.*, 2013). We have recently trialled deep brain stimulation (DBS) of the NBM as a potential therapy for both PDD and DLB (Gratwicke *et al.*, 2018, 2020*b*). This has offered the opportunity to gain unique neurophysiological insights into the function of the NBM and its cortical networks. By combining recordings of NBM local field potentials (LFP) from DBS electrodes with magnetoencephalography (MEG), we have been able to identify NBM-cortical networks that are common to both diseases (Gratwicke *et al.*, 2020*a*) including: 1) a delta/theta band (2-8 Hz) network between the NBM and temporal cortex, and a low beta band (13-22 Hz) network between the NBM and mesial motor areas. Despite this, it remains unclear how the *functional* connectivity of the NBM relates to its *structural* connectivity.

Previous work with MRI tractography reveals extensive structural projections of the NBM to cortical regions (Hepp *et al.*, 2017; Nemy *et al.*, 2020). It is likely, however, that the outputs of the basal forebrain structures are both spatially and temporally (Gielow and Zaborszky, 2017; Záborszky *et al.*, 2018) segregated representing discrete functional networks that subserve distinct roles. An improved understanding of the segregation of these networks is essential for understanding their functional and pathophysiological roles.

Here we test for spatiotemporal segregation of NBM-cortical networks, by integrating previously reported (Gratwicke *et al.*, 2020*a*) MEG and NBM LFP recordings with MRI tractography derived from open source connectomes (Horn *et al.*, 2017). Using this approach, we test for the presence of brain networks that are both *structurally* and *functionally* connected to the NBM.

## Methods

Six PDD patients and five DLB patients who participated in the clinical trials underwent combined MEG and NBM LFP recordings. Clinical characteristics of the patients and details of the surgical procedure are provided in previous clinical and MEG studies of the patients (Gratwicke *et al.*, 2018, 2020*a*, *b*). The bilateral quadripolar Medtronic leads had their two distal contacts (contacts 0 and 1) lying in the region of the NBM, and the two proximal contacts (contacts 2 and 3) in the internal segment of the globus pallidus (GPi). Study procedures were approved by the East of England Research Ethics Service Committee, and the patients gave written informed consent prior to participation.

### Electrophysiological recordings

MEG recordings were performed using a 275-channel MEG system (CTF/VSM MedTech). LFP activity recorded from DBS electrodes was collected at the same time as MEG using a BrainAmp system (Brain Products). Three bipolar channels (0-1, 1-2, 2-3) were recorded from each electrode and were high-pass filtered at 1 Hz in the hardware to avoid amplifier saturation due to large DC offsets. Rest recordings of a duration of 3 minutes were performed whilst patients were on their usual medication.

### Magnetoencephalography and tractography analysis

The aim of joint MEG and LFP analysis was to generate a Montreal Neurological Institute (MNI) space whole brain image of coherence – which we used as a measure of frequency specific *functional* connectivity – between the NBM (contact pair 0-1) or GPi (contact pair 2- and 5mm spaced grid points within the brain. For this purpose we used a single shell forward model, based on each patient’s pre-operative MRI (Oswal *et al.*, 2016), in conjunction with Dynamic Imaging of Coherent Sources (DICS) beamforming implemented in the Data Analysis in Source Space (DAiSS) toolbox for SPM12 (https://github.com/spm/DAiSS). Values at the grid points were then interpolated to produce coherence images with 2 mm resolution. Based on previous analysis (Gratwicke *et al.*, 2020*a*) we restricted our analyses of *functional* connectivity with the NBM to the delta/theta (2-8 Hz), low beta (13-22 Hz) and high beta (22-30 Hz) bands.

Lead-DBS software was used to localise electrode contacts in the aforementioned MNI space (Horn and Kühn, 2015). For the purposes of fibre tracking, a spherical region of interest centred at the midpoint of each chosen contact pair, with a radius that just encompassed each contact pair was constructed. This spherical volume was used as a seed region in an openly available group connectome (www.lead-dbs.org) which was derived from the diffusion-weighted magnetic resonance images of 32 healthy subjects within the human connectome project. Whole brain tractography fibre sets were calculated within a white-matter mask after segmentation with SPM12 using DSI studio (http://dsi-studio.labsolver.org). Fibre tracts were transformed into MNI space for visualisation. The number of fibres passing through both the spherical seed and each 2×2×2 mm cubic voxel served as an estimate of tract density and was written to a 3D image. Images corresponding to left LFP channels were flipped across the mid-sagittal plane to allow comparison of ipsilateral and contralateral sources regardless of original side. Tract density images were compared at the group level using a 2×2 ANOVA in SPM12 with factors contact location (NBM vs. GPi) and disease (PDD vs. DLB). We included covariates in order to account for both subject specific dependencies in the recordings from both hemispheres and for potential differences between recordings of the right and left sides.

### Integration of structural and functional connectivity

We used a group level voxel-wise general linear model (GLM) for determining whether structural connectivity was predictive of cortico-LFP coherence separately for each of the two frequency bands and for NBM and GPi contact pairs. After right flipping both coherence and tract density images corresponding to left LFP channels, we constructed a GLM for each voxel with tract density as the independent variable and coherence as the dependent variable. This is different from the standard SPM approach since here the design matrix is voxel-specific rather than the same for the whole brain. A cluster-based permutation test with cluster-forming threshold of p<0.01 was used to define significance. We report results significant at p<0.01 family-wise error corrected at the cluster level. F-statistics of voxels within each cluster were then written to a 3D nifty format image for visualisation.

### Directionality analysis

The effective directionality of coupling between the cortex and the NBM LFP was computed with a non-parametric variant of spectral Granger causality (Dhamala *et al.*, 2008). To determine the significance of directionality estimates, we compared the Granger estimate of original data to that of surrogate time-reversed data using a paired *t*-test (Haufe *et al.*, 2013). Taking the example of two signals A and B, with A Granger causing B, the Granger causality from A to B should be higher for the original than for the time-reversed data, giving rise to a positive difference. In contrast, the estimate of causality from B to A should be increased by time reversal thereby giving a negative difference.

### Data Availability

Exemplar code for computation of structural and functional connectivity relationships can be found on the following GitHub repository (https://github.com/AshOswal/Multimodal_Tools). Anonymised data are available from the corresponding authors on request.

## Results

### Visualisation of tracts and oscillatory networks

The upper panel of **Figure 1** displays tractography streamlines of fibres passing from the vicinity of NBM or GPi contacts to cortical structures for the PDD (turquoise) and DLB (magenta) patient groups. In the case of NBM contacts there are extensive fibre connections to numerous cortical regions (‘corticopetal’ connections). MEG derived cortical networks exhibiting coherence with both the NBM and GPi at delta/theta (2-8 Hz; red), low beta (13-22 Hz; yellow) and high beta (22-30 Hz; blue) frequencies are also displayed. Note that the MEG networks were derived after right flipping of images corresponding to the left LFP and are therefore predominantly ipsilateral. The lower panel in **Figure 1** displays the results of the 2×2 ANOVA comparing tract densities with factors contact location (NBM vs. GPi) and disease (PDD vs. DLB). The only significant finding was a main effect of contact location, such that NBM contacts displayed greater *structural* connectivity with a region including the hippocampus (blue contour), the lingual gyrus (turquoise contour), the calcarine cortex (green contour), and occipital cortex (magenta contour). Due to the lack of a main effect of disease for both *structural* and *functional* connectivity, we pooled across the disease states for further analysis, separately for GPi and NBM contacts.

**Figure 1.**
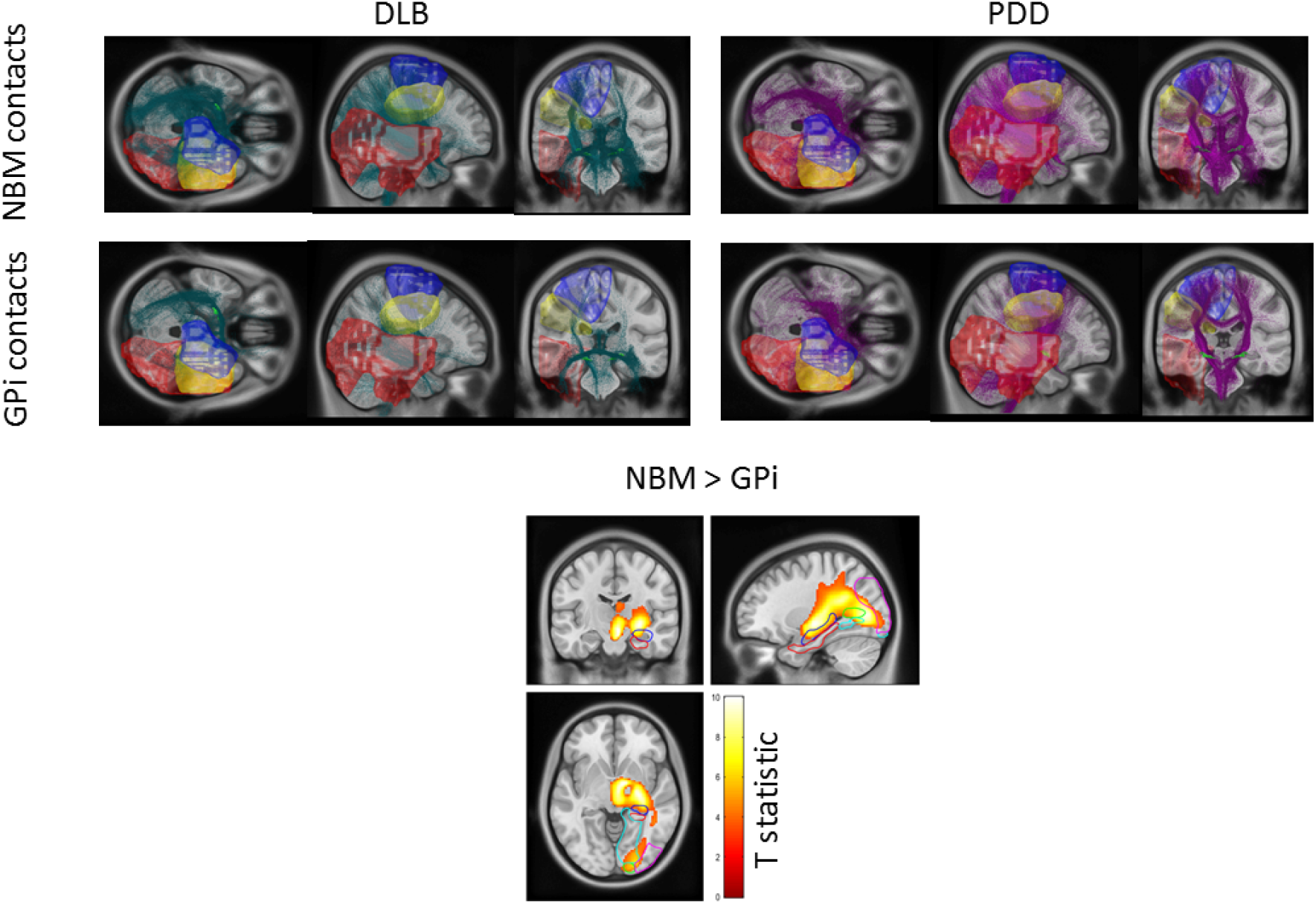
Visualisation of tracts and oscillatory networks. **Upper panel:** fibre streamlines passing from the vicinity of NBM and GPi contacts to cortical regions are displayed for the two disease groups on a T1-weighted MRI scan. For DLB patients, fibres are coloured turquoise, whilst for PDD patients they are coloured magenta. MEG-derived cortical networks displaying coherence with both the GPi and NBM in the delta/theta (red surface), low beta (blue surface) and high beta (yellow surface) bands are also displayed. **Lower panel**: Statistical Parametric Map (SPM) displaying the results of the 2×2 ANOVA with factors disease (PDD vs. DLB) and contact location (NBM vs. GPi). The statistical T-image displays regions that have significantly greater structural connectivity with the NBM than with the GPi (main effect of location). These include the hippocampus (blue contour), lingual gyrus (turquoise contour), calcarine cortex (green contour), and occipital cortex (magenta contour). The Parahippocampal gyrus is indicated by the red contour.

### Structural connectivity correlates with functional connectivity within spatially and spectrally distinct NBM-cortical networks

Panel (A) in **Figure 2** reveals cortical regions where NBM-cortical tract density was predictive of NBM-cortical coherence in the low beta frequency range (13-22 Hz). The significant cluster encompasses the SMA. Panel (B) in contrast reveals regions where NBM-cortical tract density was predictive of NBM-cortical coherence in the delta/theta frequency range (2-8 Hz). Three clusters are displayed that include the parahippocampal gyrus (red contour), the inferior temporal cortex and the lingual gyrus (turquoise contour). Source extracted coherence spectra for the peak locations in panels (A) and (B) are shown in the panels of **Figure 3A**. These reveal peaks in the beta and delta/theta bands respectively.

**Figure 2.**
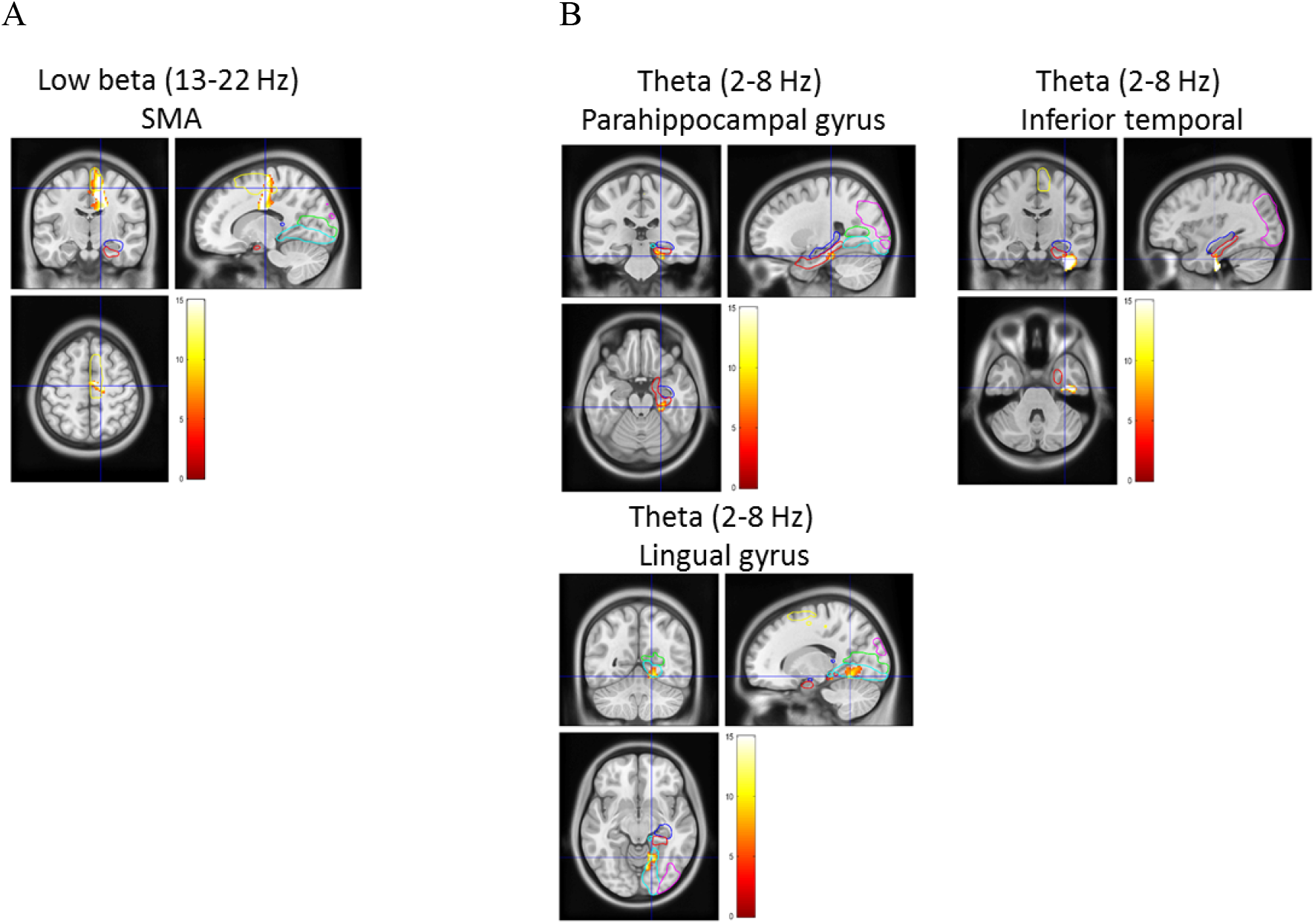
Spatially and spectrally distinct cortico-NBM networks with correlated structural and functional connectivity. **(A):** a statistical image showing premotor areas (including SMA; yellow contour) where structural connectivity and coherence with the NBM in the low beta band are correlated. **(B):** regions where structural connectivity and coherence with the NBM in the delta/theta band are correlated. Three clusters are displayed that include the parahippocampal gyrus, the inferior temporal cortex and the lingual gyrus. Images are superimposed on a T1-weighted MRI scan and the colourbar represents the value of the F-statistic. Contours of the supplementary motor area (SMA; yellow contour) hippocampus (blue), parahippocampal gyrus (red), lingual gyrus (turquoise contour), calcarine cortex (green), and occipital cortex (magenta contour) are also shown.

**Figure 3.**
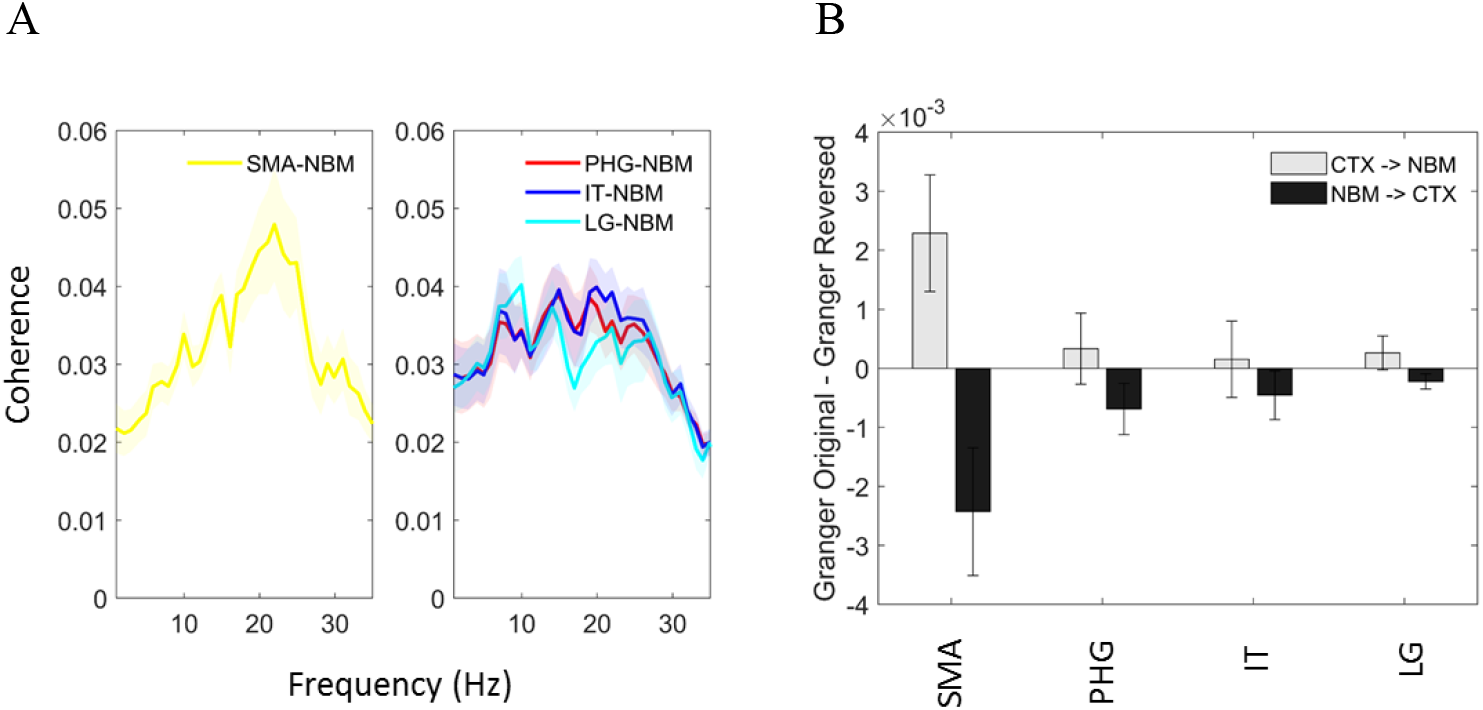
Group spectra and directionality analysis. **(A):** left panel shows coherence spectra computed between the NBM LFP and the location of the peak F statistic of the correlation between beta band coherence and tract density which was within the SMA. Similarly, in the right panel coherence between the NBM LFP and cortical locations displaying a correlation between structural and functional connectivity in the delta/theta band is plotted (PHG = parahippocampal gyrus, IT = inferior temporal cortex, LG = lingual gyrus). **(B):** Group mean differences in Granger causality between the original data and time reversed data are averaged across the beta band for the SMA and across the delta/theta band for the PHG, the IT and LG. The difference in Granger causality is significantly greater than zero in the direction of SMA leading the NBM in the beta band.

Importantly we have observed differences in the *structural* connectivity profiles of the NBM and GPi, despite their *functional* connectivity (Gratwicke *et al.*, 2020*a*) profiles being similar. Furthermore, no significant relationship between GPi-cortical tract density and GPi-cortical coherence was observed in any of the three frequency bands.

### Directionality of coupling within structurally and functionally connected cortico-NBM networks

For the purposes of directionality analysis, we derived source time series from the locations of the peak F-statistics of the correlation between structural and functional connectivity (see **Figure 2**), separately for the delta/theta and beta bands. For the SMA-NBM beta band network, the difference in granger causality, averaged across the beta band, was significantly greater than zero in the direction of SMA leading the NBM (F_(1,10)_ = 5.63, p = 0.039; **Figure 3B**) suggesting cortical driving of the NBM. In contrast, for the parahippocampal gyrus-NBM (cortex leading F_(1,10)_ = 0.21, p = 0.66; NBM leading F_(1,10)_ = 2.63, p = 0.14), the inferior temporal-NBM (cortex leading F_(1,10)_ = 0.03, p = 0.86; NBM leading F_(1,10)_ = 1.2, p = 0.30) and the lingual gyrus-NBM networks (cortex leading F_(1,10)_ = 1.17, p = 0.31; NBM leading F_(1,10)_ = 2.67, p = 0.13), the difference in granger causality averaged across the delta/theta band was not significant in any particular direction, hence suggesting bidirectional patterns of communication.

## Discussion

In this report we develop a methodology to explore the intersection of *structural* and *functional* connectivity and using this we identify distinct brain networks that are both structurally and functionally connected to the NBM in patients with DLB and PDD. Importantly the correlation of *structural* and *functional* connectivity was specific to the NBM rather than to the GPi, which we hypothesise may be reflective of a monosynaptic input-output relationship between the NBM and cortical structures (Gielow and Zaborszky, 2017; Záborszky *et al.*, 2018). The GPi in contrast has polysynaptic cortical connections via intermediate structures (Nambu, 2007) such as the thalamus and other basal ganglia nodes.

We identified three spatially and spectrally distinct NBM-cortical networks with overlapping structural and functional connectivity. Firstly we identified a low beta band network (13-22 Hz) between the NBM and the SMA, an area known to be important in the volitional control of movement (Nachev *et al.*, 2008). The existence of this network is consistent with previous tracer studies in humans, rodents and primates demonstrating monosynaptic connections projecting from both the NBM to the sensorimotor areas and vice-versa (Mesulam and Mufson, 1984; Mesulam and Geula, 1988; Gielow and Zaborszky, 2017). It is hypothesised that such cortico-NBM projections may play important roles in motor plasticity and skill learning. Importantly, within this network we observed that cortical activity tended to drive NBM activity. This direction of information flow appears more consistent with the hypothesised ‘top-down’ model of the fronto-parietal attention network, wherein direct frontal cortical connections to NBM modulate its cholinergic output to other cortical areas to amplify processing of attention demanding signals (Duque *et al.*, 2000; Sarter *et al.*, 2005). Furthermore, modulation of attention within this network has been shown to be related to functional connectivity in the beta band (Buschman and Miller, 2007). Interestingly, in addition to driving NBM activity at low beta frequencies, the SMA also couples with and drives subthalamic nucleus (STN) activity at high beta (21-30 Hz) band frequencies which are likely to be reflective of hyperdirect pathway activity (Oswal *et al.*, 2016). These findings therefore indicate that the outputs of the SMA to different anatomical structures may be spectrally segregated.

Secondly, two spatially distinct delta/theta (2-8 Hz) band networks were seen; one including inferior and mesial temporal lobe structures such as the parahippocampal gyrus and a second including lingual gyrus. Based on spatial location, it is likely that these networks may have important roles in memory and visual function respectively (Gratwicke *et al.*, 2013; Ballinger *et al.*, 2016; Huppé-Gourgues *et al.*, 2018). In contrast to the beta band network, we did not identify a net directionality of information flow within the theta band network. It is possible that this is likely to reflect bidirectional patterns of communication between the NBM and mesial temporal structures such as the hippocampus and parahippocampal cortex, which have also been demonstrated with tracer studies (Mesulam and Geula, 1988; Gratwicke *et al.*, 2013).

Our findings should be interpreted in light of the limitation that we used tractography data from normative connectomes rather than from individual subjects. Nevertheless, this approach also offers a major advantage in that connectome data are acquired with specialised hardware and large cohort sizes leading to connectivity estimates with improved signal-to-noise ratio compared to what would be possible with individual patient data. Connectome data have been successfully leveraged recently to study the mechanisms of DBS action (Horn *et al.*, 2017) and also to explore how data from disparate lesion studies can be integrated to understand the role of brain networks in disease (Fox, 2018).

In summary, our findings demonstrate for the first time a relationship between structural and functional connectivity within the cortico-NBM network and motivate future studies aimed at disentangling the relative contribution of the identified networks in both normal neurophysiological functioning and neurological disease.

## Acknowledgements

AO is supported by an NIHR Academic Clinical Lectureship and an Academy of Medical Sciences Starter Grant. The Wellcome Centre for Human Neuroimaging is supported by core funding from Wellcome [203147/Z/16/Z]. The work was supported by the UK MEG community Medical Research Council grant MK/K005464/1. PB is supported by the Medical Research Council (MC_UU_12024/5). Laszlo Zaborszky is supported by the NIH/NINDS grant number: 2RF1NS023945-28. TF receives funding from the Brain Research Trust, Michael J Fox Foundation, European Union FP-7 and John Black Charitable Foundation. Ludvic Zrinzo and HA are supported by a grant from the Brain Research Trust (157806) and the National Institute for Health Research University College London Hospitals Biomedical Research Centre. The Unit of Functional Neurosurgery is supported by the Parkinson's Appeal and the Sainsbury Monument Trust.

